# Structural basis for direct NGF/TrkA blockade by an analgesic antibody

**DOI:** 10.64898/2026.06.30.735605

**Authors:** Harsh Bansia, Elisa Damo, Eliezra Glasser, Renato Bruni, Shohei Koide, Nigel W. Bunnett, Amédée des Georges

## Abstract

The NGF/TrkA signaling axis is a central mediator of inflammatory and chronic pain, where injury-induced NGF binds and activates TrkA on nociceptive neurons to drive peripheral sensitization and persistent pain states. Despite its therapeutic promise, targeting this pathway is limited by adverse effects of systemic NGF sequestration such as rapidly progressive osteoarthritis and poor isoform selectivity of Trk kinase inhibitors leading to off-target neurological effects. Targeting the TrkA extracellular domain (TrkA_ECD_) offers a pathway to achieve high isoform selectivity while avoiding these complications. However, the precise structural basis for selective TrkA neutralization remains poorly understood. Monoclonal antibody (mAb) 42F5-15 inhibits TrkA-mediated signaling and increases pain threshold. Here, we report the high-resolution (2.60 Å) cryo-EM structure of the TrkA_ECD_ in complex with the Fab region of the TrkA-neutralizing mAb 42F5-15. Structural analysis reveals that the antibody epitope overlaps the NGF-binding interface, consistent with orthosteric inhibition and distinct from previously proposed allosteric mechanisms. The epitope includes residues conserved in TrkA but divergent in TrkB and TrkC, providing a structural basis for receptor isoform selectivity. Furthermore, we demonstrate *in vivo* that the mAb 42F5-15 potently mitigates mechanical allodynia and nociceptive sensitization. These findings establish a structural framework for the development of selective extracellular TrkA-targeted therapies for safer, non-opioid chronic pain management.

## Introduction

Nerve Growth Factor (NGF) and its high-affinity receptor tropomyosin receptor kinase A (TrkA), carry out pleiotropic roles across the lifespan, shifting from essential embryonic functions in neuronal survival and innervation [1–3] to a critical mediator of adult nociceptive pain [4,5]. NGF is rapidly upregulated at sites of tissue injury and inflammation [6], where it binds and activates TrkA [7] on peripheral nociceptive sensory neurons. The formation of active NGF/TrkA complex triggers downstream signaling cascades [8] that drive rapid peripheral sensitization - lowering the activation thresholds for thermal, mechanical, and chemical stimuli [9,10] - and induce transcriptional changes that promote and maintain chronic pain states [11–13]. The clinical relevance of the NGF/TrkA axis as a therapeutic target for chronic pain [14,15] is supported by observations from Congenital Insensitivity to Pain with Anhidrosis (CIPA), a rare hereditary disorder [16]. CIPA is caused by loss-of-function mutations in the *NTRK1* gene, which encodes the TrkA receptor, leading to a profound inability to perceive painful stimuli [16] and thereby demonstrating the central role of this pathway in human nociception [14,15,17].

Despite the therapeutic potential of targeting the NGF/TrkA axis [15,18], existing strategies face significant limitations. Systemic NGF sequestration with anti-NGF monoclonal antibodies such as *tanezumab* [19] and *fasinumab* [20] effectively reduces pain [21] by preventing NGF from activating TrkA signaling, but has been associated with serious adverse events, including rapidly progressive osteoarthritis (RPOA) [22]. Although the underlying mechanism remains incompletely understood, broad and non-selective inhibition of NGF signaling may also disrupt p75 neurotrophin receptor (p75NTR)-mediated pathways [23,24], which have been implicated in skeletal cell migration and bone repair [25]. Conversely, small-molecule inhibitors targeting the intracellular TrkA kinase domain [26,27] often suffer from limited isoform selectivity due to the high structural conservation of the ATP-binding pockets across the TrkA, TrkB, and TrkC kinase domains. As a result, off-target inhibition of TrkB and TrkC may produce undesirable neurological effects, given their essential roles in synaptic plasticity, cognition, neuronal survival, and motor function [28].

The TrkA extracellular domain (TrkA_ECD_), particularly the second immunoglobulin-like (IgC2) domain, contributes significantly to NGF binding affinity and isoform selectivity among Trk receptor isoforms [29,30]. Accordingly, targeting the TrkA_ECD_ has been proposed as a strategy to selectively prevent NGF-induced TrkA activation without affecting p75NTR-mediated signaling. This approach may, in principle, avoid the limitations associated with systemic NGF sequestration and the off-target effects of small-molecule Trk kinase domain inhibitors. Consequently, TrkA_ECD_ targeting has emerged as a promising alternative strategy to achieve higher isoform selectivity while potentially reducing systemic adverse effects. Although previous studies, most notably involving the antibody MNAC13, have demonstrated the feasibility of neutralizing NGF/TrkA signaling axis by targeting TrkA_ECD_ [31,32], the precise structural basis of this mechanism remains poorly understood. Recently, antibody 42F5-15 has been shown to achieve higher affinity, better specificity and stronger activity and its use is proposed as a strategy for developing drugs to treat pain including inflammatory, neuropathic, cancer and osteoarthritis [33]. However, the absence of high-resolution structures of anti-TrkA antibodies in complex with the TrkA_ECD_ has limited mechanistic insight into how these molecules inhibit NGF/TrkA signaling.

Here, we address this knowledge gap by reporting the high-resolution (2.60 Å) cryo-EM structure of the TrkA_ECD_ in complex with a TrkA-neutralizing monoclonal antibody, 42F5-15. Our structural analysis reveals that the epitope of mAb 42F5-15 overlaps with the NGF-binding interface on TrkA_ECD_ Ig-C2 domain, consistent with an orthosteric mode of inhibition that is mechanistically distinct from previously proposed allosteric mechanisms for MNAC13. Furthermore, we identify epitope residues that are conserved in TrkA but divergent in TrkB and TrkC, thus establishing structural basis for Trk receptor isoform selectivity. Finally, functional *in-vivo* assays demonstrate that the mAb 42F5-15 effectively mitigates mechanical allodynia and reduces nociceptive sensitization. Collectively, this study provides structural insights into selective extracellular targeting of TrkA_ECD_, elucidating mechanistic details of the strategy to overcome systemic limitations associated with current NGF/TrkA-targeted therapies and supporting the development of more precise non-opioid approaches for chronic pain management.

## Results

### 1. Cryo-EM Structure of the TrkA Ectodomain Complex with Fab Region of Monoclonal Antibody 42F5-15

To minimize conformational heterogeneity and improve particle alignment for cryo-electron microscopy (cryo-EM), Fab regions were generated from the monoclonal antibody (mAb) 42F5-15. Fab regions were specifically chosen to eliminate the rotational dynamicity and flexibility inherent in the bivalent IgG hinge region, which can otherwise impede high-resolution 3D reconstruction by introducing structural noise during particle alignment. Cryo-EM grid preparation, data collection, data processing, model building and refinement are described in **Methods**.

The cryo-EM analysis yielded distinct 3D reconstructions that together provide a multi-scale view of the TrkA_ECD_/42F5-15_Fab_ complex **(Fig. 1)**. TrkA_ECD_ consists of leucine-rich repeat (LRR) and cysteine-rich regions, followed by two immunoglobulin-like domains, classified as Ig-C1 and Ig-C2, respectively **(Fig. 1d).** Among these, the Ig-C2 domain plays a central role in neurotrophin binding, with structural and mutagenesis studies identifying it as a key determinant of NGF interaction with TrkA [30]. A consensus map was resolved at a global resolution of 2.90 Å based on gold-standard Fourier shell correlation of 0.143 criterion (**Fig. 1a**), exhibiting density for the TrkA ECD, including LRR, Ig-C1 and Ig-C2 domains **(Fig. 1e)**. This map allowed for the unambiguous placement of the TrkA ECD relative to the Fab **(Fig. 1e)**. The 2.90 Å map reveals that the TrkA_ECD_/42F5-15_Fab_ complex adopts a distinct, elongated architecture reminiscent of a “***scorpion-like***” silhouette (**Fig. 1a, e**). In this arrangement, the Fab region serves as the stable, “body” at the base of the complex (**Fig. 1e**). The TrkA ECD, anchored at the paratope, curves upward, forming the flexible “tail” (**Fig. 1e**). However, lower local resolution in the LRR domain suggested significant conformational flexibility in the peripheral domains of TrkA. Separate reconstructions from different sets of particles with partial TrkA ECD achieved higher resolutions of 2.70 Å (**Fig. 1b**) and 2.60 Å (**Fig. 1c**). Whereas both reconstructions exhibited well-defined density for the Fab and the primary interacting domain, Ig-C2 of TrkA, the peripheral TrkA domains, LRR in the 2.70 Å map and LRR, Ig-C1 in the 2.60 Å map, remained unresolved **(Fig. 1b, c)**. The absence of these peripheral domains in the high-resolution maps may be due to conformational flexibility, partial proteolytic cleavage, or a combination of both.

**Figure 1:**
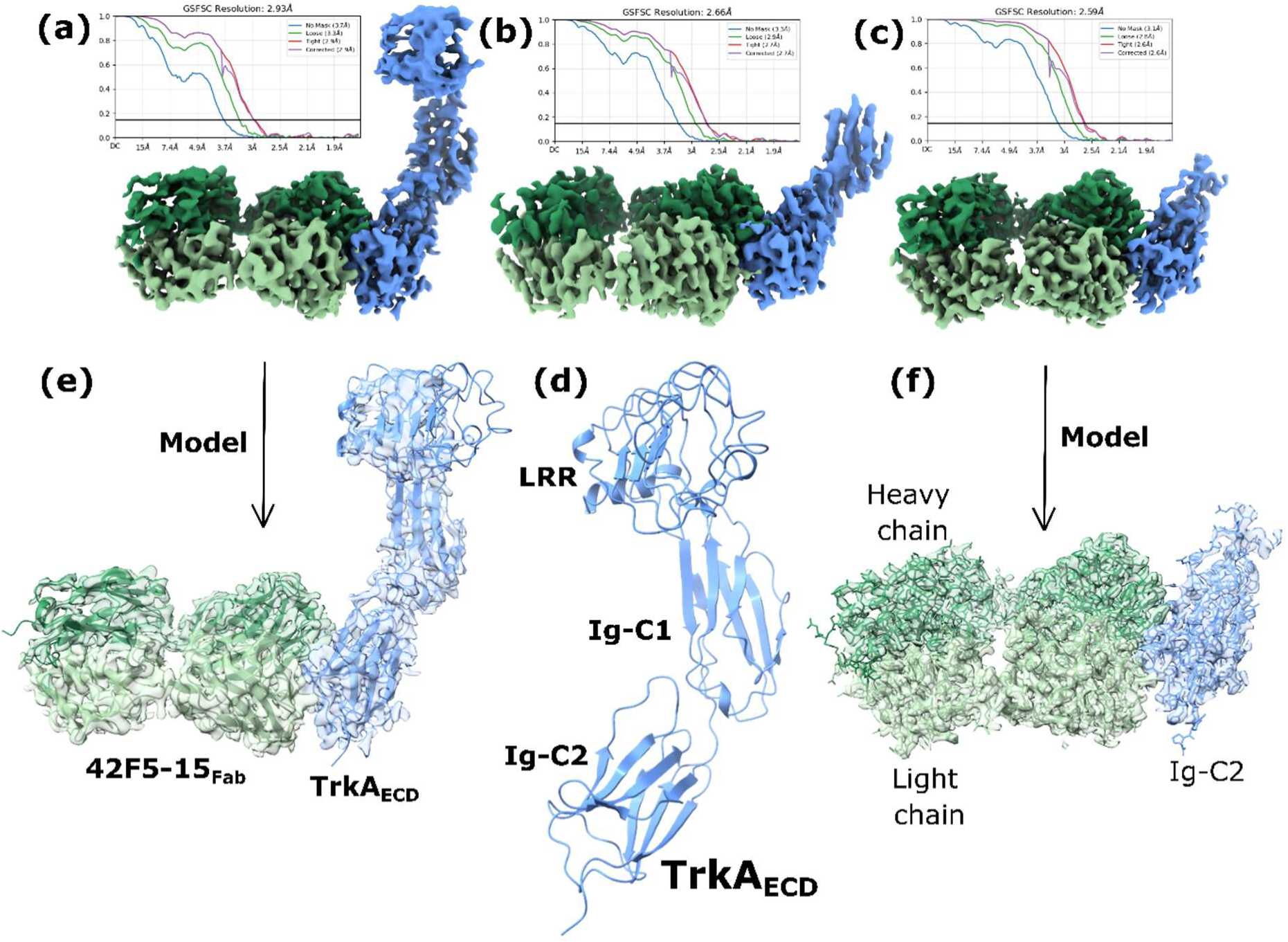
Cryo-EM structure of the TrkA_ECD_/42F5-15_Fab_ complex. Distinct 3D reconstructions at resolutions **(a)** 2.90 Å, **(b)** 2.70 Å, and **(c)** 2.60 Å, based on gold-standard Fourier shell correlation of 0.143 criterion, provide a multi-scale view of the TrkA_ECD_/42F5-15_Fab_ complex. mAb 42F5-15_Fab_ (Green/pale-green; heavy/light chain); TrkA_ECD_ (sky-blue). **(d)** Cartoon representation showing leucine-rich repeat (LRR) and cysteine-rich regions, and two immunoglobulin-like domains, classified as Ig-C1 and Ig-C2, of TrkA_ECD_. **(e)** The 2.90 Å map captures the complete molecular envelope of TrkA_ECD_/42F5-15_Fab_ complex, and **(f)** the 2.60 Å reconstruction reveals side-chain densities to model specific molecular interactions at the paratope-epitope interface. **(e, f)** Transparent cryo-EM density of the maps overlayed with the models of TrkA_ECD_/42F5-15_Fab_ complex to show the map-model fit. Mapping of TrkA LRR, Ig-C1 and Ig-C2 domains of in the TrkA_ECD_/42F5-15_Fab_ complex reveals that the mAb 42F5-15 binds to TrkA Ig-C2 domain, which is also the NGF binding site.

### 2. Structural Analysis of the TrkA_ECD_/42F5-15_Fab_ Interface and Epitope Mapping

The 2.90 Å and 2.60 Å maps were utilized for atomic model building and analysis of TrkA_ECD_/42F5-15_Fab_ complex. While the density from the 2.90 Å reconstruction provided the necessary placement and connectivity for the TrkA LRR, Ig-C1 and Ig-C2 domains **(Fig 1e)**, the resolvability in the 2.60 Å reconstruction enabled modeling at the side-chain level (**Fig. 1f**) to elucidate specific molecular interactions at the paratope-epitope interface (**Fig. 2a**). The interaction interface of the TrkA_ECD_/42F5-15_Fab_ complex buries a total surface area of approximately 1,565 A^2^, comprising 793.7 A^2^ of TrkA and 771.6 A^2^ of the Fab. These buried regions represent 4 % of the total solvent-accessible surface area of the TrkA (19,772.7 A^2^) and 6.7 % of the Fab (11,727.5 A^2^), respectively. This interface footprint is consistent with high-affinity antibody-antigen complexes [34].

**Figure 2.**
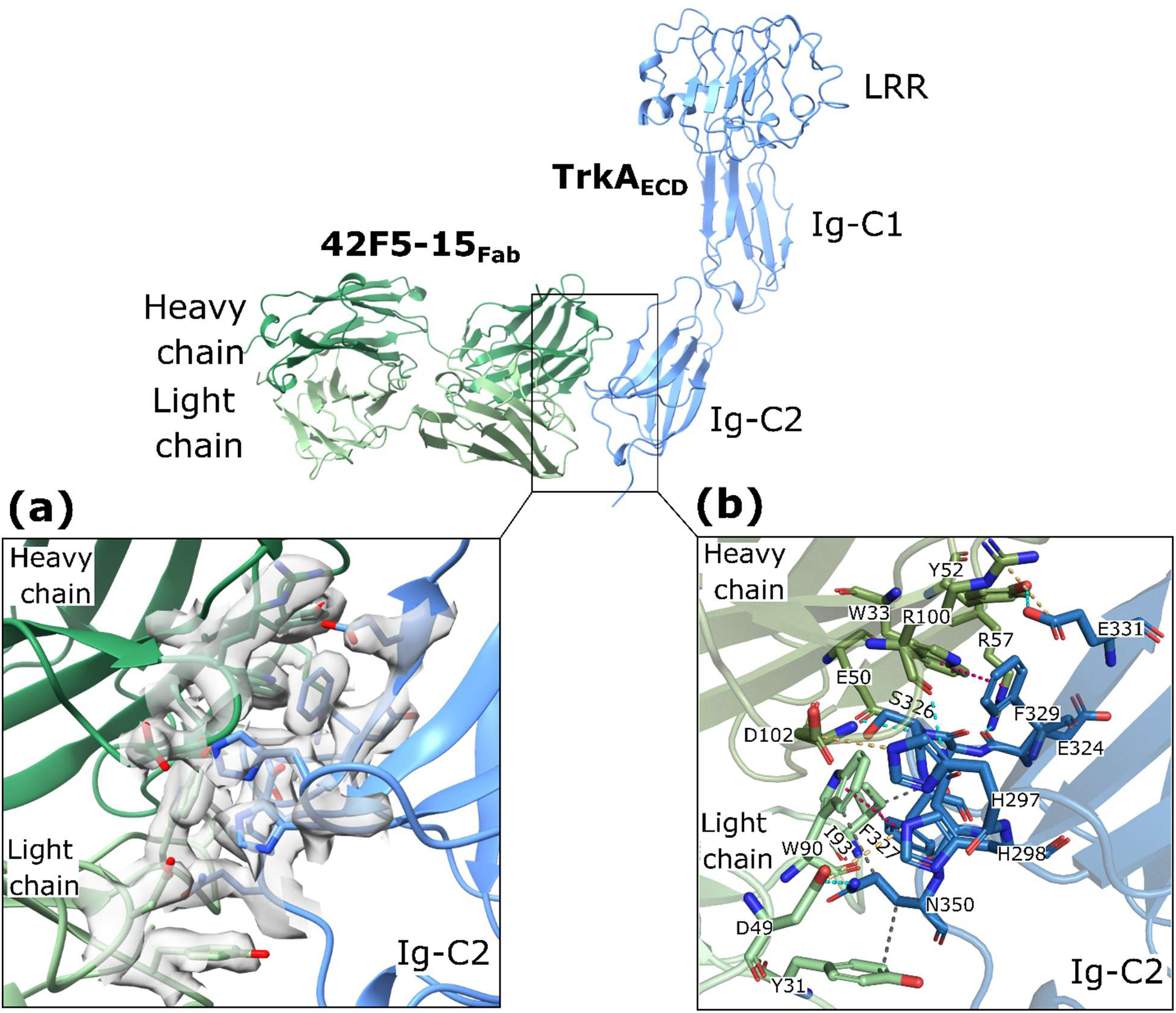
Molecular interactions at the TrkA_ECD_/42F5-15_Fab_ complex. **(a)** Interface of the TrkA_ECD_/42F5-15_Fab_ model (cartoon representation) is overlayed with the cryo-EM density at the interface to show that the modeling of the side-chains (sticks representation) and calculation of specific molecular interactions at the binding interface is guided by side-chain density definitions (transparent gray surface) revealed in the high-resolution 2.60 Å reconstruction. **(b)** Analysis of TrkA_ECD_/42F5-15_Fab_ interface reveals that the mAb 42F5-15 (Green/pale-green; heavy/light chain) recognizes a conformational epitope on TrkA Ig-C2 domain (sky-blue) and its paratope comprises of CDRs H2, H3, L1, and L2. Epitope-paratope residues are shown as stick model colored as per the color convention for TrkA_ECD_/42F5-15_Fab_ model. Molecular interactions at the TrkA_ECD_/42F5-15_Fab_ interface are shown as dashed lines: Hydrogen bonds (cyan), electrostatic/ionic (orange), hydrophobic (dark-gray) and pi-stacking (magenta).

Analysis of the TrkA_ECD_/42F5-15_Fab_ complex structure reveals that the mAb 42F5-15 recognizes a conformational epitope on the Ig-C2 domain of TrkA. The epitope on TrkA Ig-C2 comprises of residues H297, H298, N323, E324, T325, S326, F327, F329, T330, E331, F332, R347, N349, Q350, P351, T352 and M379. The epitope on TrkA Ig-C2 is recognized by a paratope on mAb 42F5-15 through a combination of hydrogen bonding, electrostatic and hydrophobic interactions. Residues E50, Y52, R57 of CDR H2, R100, Y101 of CDR H3, Y31, H33 of CDR L1 and D49 of CDR L2 form hydrogen bonds with residues H298, E324, S326, E331, R347, N349, Q350, P351 of TrkA Ig-C2. Residues R100, D102 of CDR H3 and D49 of CDR L2 form electrostatic/salt-bridge interactions with residues H297, H298 and E331of TrkA Ig-C2. Residues W33 of CDR H1 and W90 of CDR L3 form pi stacking interactions with residues F327 and F329 of TrkA Ig-C2. Residues W90, I93 of CDR L3 and Y31 of CDR L1 form hydrophobic interactions with residues T325, F327, N349, Q350 of TrkA Ig-C2. Details of the interacting residues, as described above for the interface of TrkA_ECD_/42F5-15_Fab_ complex, are illustrated in **Figure 2b**.

### 3. Computational Mapping of MNAC13 Epitope

MNAC13 is an extensively characterized anti-TrkA monoclonal antibody with established *in vitro* and *in vivo* neutralizing efficacy against the NGF/TrkA axis, delivering potent analgesia in inflammatory pain models and sustained efficacy in neuropathic pain [31]. Previous epitope mapping localized the MNAC13 binding site to residues (AGWILTELEQSATVMKS) within the TrkA Ig-C1 domain [32]. Although this region is part of a linear sequence, the epitope is predominantly conformational, as evidenced by phage display “mimotopes” that lack sequence homology with TrkA Ig-C1 domain but share critical biochemical features, such as a hydrophobic patch followed by acidic residues [32]. Despite the availability of the MNAC13 Fab crystal structure [32], the absence of a TrkA_ECD_/MNAC13_Fab_ complex structure limits understanding of molecular interactions at the interface and precise neutralization mechanism of MNAC13. To address this, AlphaFold3 [35] was leveraged to predict the TrkA_ECD_/MNAC13_Fab_ complex structure (**Fig. 3a**), utilizing TrkA_ECD_ sequence from UniProt (P04629) and the MNAC13_Fab_ sequence from MNAC13_Fab_ crystal structure (PDB 1SEQ), respectively. The resulting models, filtered based on established epitope mapping and phage display data [32], consistently localized the binding interface to the Ig-C1 domain of TrkA (around residues AGWILTELEQSATVMKS) (**Fig. 3b, c**) rather than the TrkA Ig-C2 domain which is the primary NGF binding site and molecular insights into the TrkA_ECD_/MNAC13_Fab_ complex interface.

**Figure 3.**
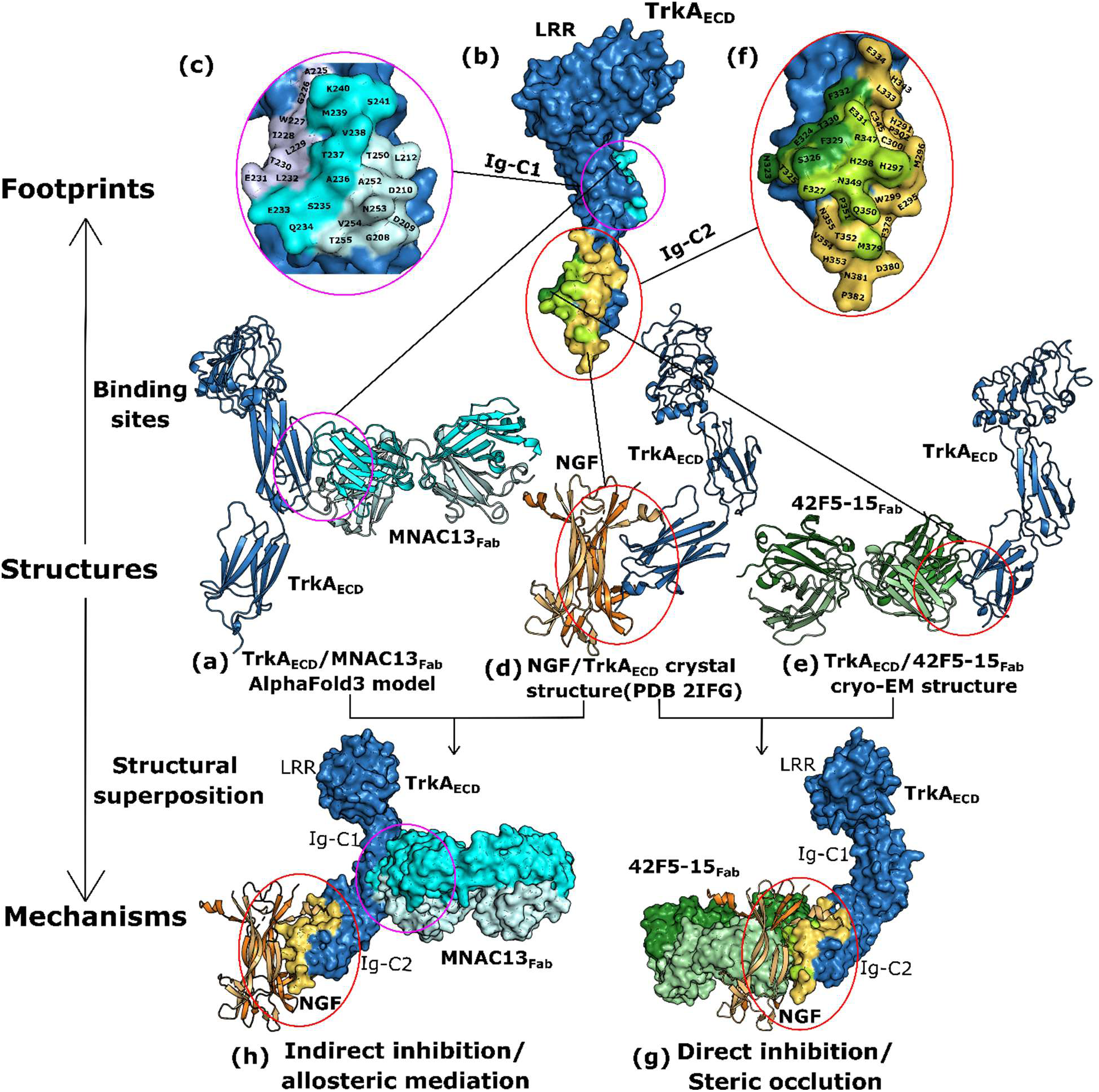
Structural basis for NGF/TrkA blockade by mAb 42F5-15 and its distinct TrkA antagonism relative to mAb MNAC13. **(a)** TrkA_ECD_/MNAC13_Fab_ AlphaFold3 model (this study) was used to map the MNAC13_Fab_ (cyan/pale-blue; heavy/light chains) footprints on **(b)** TrkA_ECD_ (surface representation, sky-blue) Ig-C1 domain. **(c)** TrkA Ig-C1 residues unique to MNAC13_Fab_ binding as inferred from the AlphaFold3 model, MNAC13_Fab_ binding as inferred from epitope mapping studies and common to AlphaFold3 model and epitope mapping, are labeled on the TrkA IgC1 domain (surface representation, colored pale-blue for AlphaFold3, light-blue for epitope mapping studies and cyan for residues common to AlphaFold3 model and epitope mapping, respectively). **(d)** NGF/TrkA_ECD_ crystal structure (PDB 2IFG, only TrkA monomer is shown for comparison) and **(e)** TrkA_ECD_/42F5-15_Fab_ cryo-EM structure (this study) were used to map the 42F5-15_Fab_ (green/pale-green; heavy/light chains) and NGF (yellow/orange dimer) footprints on **(b)** TrkA_ECD_ (surface representation, sky-blue) Ig-C2 domain. **(f)** TrkA Ig-C2 residues unique to 42F5-15_Fab_ binding, NGF binding and common to binding both, 42F5-15_Fab_ and NGF, are labeled on the TrkA IgC2 domain (surface representation, colored green for 42F5-15_Fab_, yellow for NGF and lime for shared/overlapping 42F5-15_Fab_ and NGF binding, respectively). Structural superposition of TrkA_ECD_/42F5-15_Fab_ cryo-EM structure, TrkA_ECD_/MNAC13_Fab_ AlphaFold3 model and NGF/TrkA_ECD_ crystal structure through the TrkA_ECD_ reveals that the **(g)** mAb 42F5-15 sterically occludes NGF binding and directly interferes with the formation of the NGF/TrkA complex whereas **(h)** the mAb MNAC13 does not directly compete with NGF for TrkA binding. NGF is shown as cartoon representation to represent its steric occlusion by mAb 42F5-15 (surface representation, cutting into NGF cartoon).

### 4. Comparative Structural Insights into TrkA Antagonism by mAbs 42F4-15 and MNAC13

Structural and mutagenesis studies have identified the C-terminal Ig-C2 domain of TrkA as the primary ligand-binding site for NGF [7,29,30]. Comparison of the NGF/TrkA crystal structure (PDB 2IFG) (**Fig 3d**) with our cryo-EM structure of TrkA_ECD_/42F5-15_Fab_ complex (**Fig. 3e**) revealed that NGF and 42F5-15 bind overlapping epitopes **(Fig. 3b)**. TrkA residues H297, H298, T325, F327, E331, R347, N349, Q350, P351, T352 and M379 are common to TrkA/NGF and TrkA_ECD_/42F5-15_Fab_ interfaces (**Fig. 3f**). In the NGF/TrkA crystal structure, residues E11, I31, H84 and R103 of NGF hydrogen bonds with residue H297, R347, T352, Q350 and N349 of TrkA while residue E11 of NGF forms a salt bridge with R347 of TrkA [7,30]. TrkA Ig-C2 residues which interact with NGF namely, H297, R347, T352, Q350 and N349, also interact with mAb 42F5-15 residues through hydrogen bonding and electrostatic interactions as described above (**Fig. 2b**).

Thus, the mAb 42F5-15 competes for the TrkA residues that are essential molecular determinants for its interaction with NGF (**Fig. 3f**). The structural superposition further revealed that the mAb 42F5-15 paratope sterically occludes the interface and directly interferes with the formation of the NGF/TrkA complex (**Fig. 3g**). In contrast, structural superposition of the predicted TrkA_ECD_/MNAC13_Fab_ model with the NGF/TrkA crystal structure through the TrkA_ECD_ reveals that the MNAC13 footprint is sequestered within the TrkA Ig-C1 domain, which is far removed from the NGF binding hotspot localized to the TrkA Ig-C2 domain (**Fig. 3b, c**). This indicates that MNAC13 is not structurally positioned to interfere with the initial docking of NGF on TrkA (**Fig. 3h**).

This analysis demonstrates a fundamental mechanistic distinction between mAbs 42F5-15 and MNAC13 based on their respective binding sites on the TrkA ectodomain (**Fig. 3b**). While MNAC13 is recognized as a potent neutralizing antibody, its footprint on the TrkA Ig-C1 domain is distal to the primary NGF binding site (**Fig. 3b**). Consequently, NGF binding to TrkA ECD is not inhibited by MNAC13 *in vitro* [32] and, its ability to inhibit TrkA activation by NGF *in vivo* likely arises through an indirect mechanism, such as interfering with conformational transitions required for functional receptor activation [32]. In contrast, 42F5-15 functions as a high-affinity, direct competitive antagonist of NGF that prevents the formation of NGF/TrkA signaling complex through direct structural overlap (**Fig. 3h**).

### 5. Structural Basis for Species-dependent TrkA Affinity and Trk Isoform Selectivity of the mAb 42F5-15

To evaluate the cross-species reactivity of the antibody between human and mouse TrkA orthologs, the binding affinity of the mAb 42F5-15 was measured for both. The measurements revealed high affinity binding to human TrkA (*K_d_* = 0.5 nM), whereas affinity for mouse TrkA was significantly reduced (*K_d_* = 39 nM) (**Fig. 4a**). This 78-fold decrease in affinity suggests that the sequence variations between the two species at the epitope site (TrkA Ig-C2 domain) may have a profound impact on the stability of the antibody-antigen complex. Structure-based sequence alignment of the Ig-C2 domains of human and mouse TrkA revealed that the mAb 42F5-15 epitope is identical between both species except for a single residue (**Fig. 4b**). Specifically, human TrkA contains a Glutamate at position 331 (E331), which is substituted by a Glutamine (Q333) at the structurally equivalent position in mouse TrkA (**Fig. 4b**) and this substitution may contribute to the observed affinity difference.

**Figure 4.**
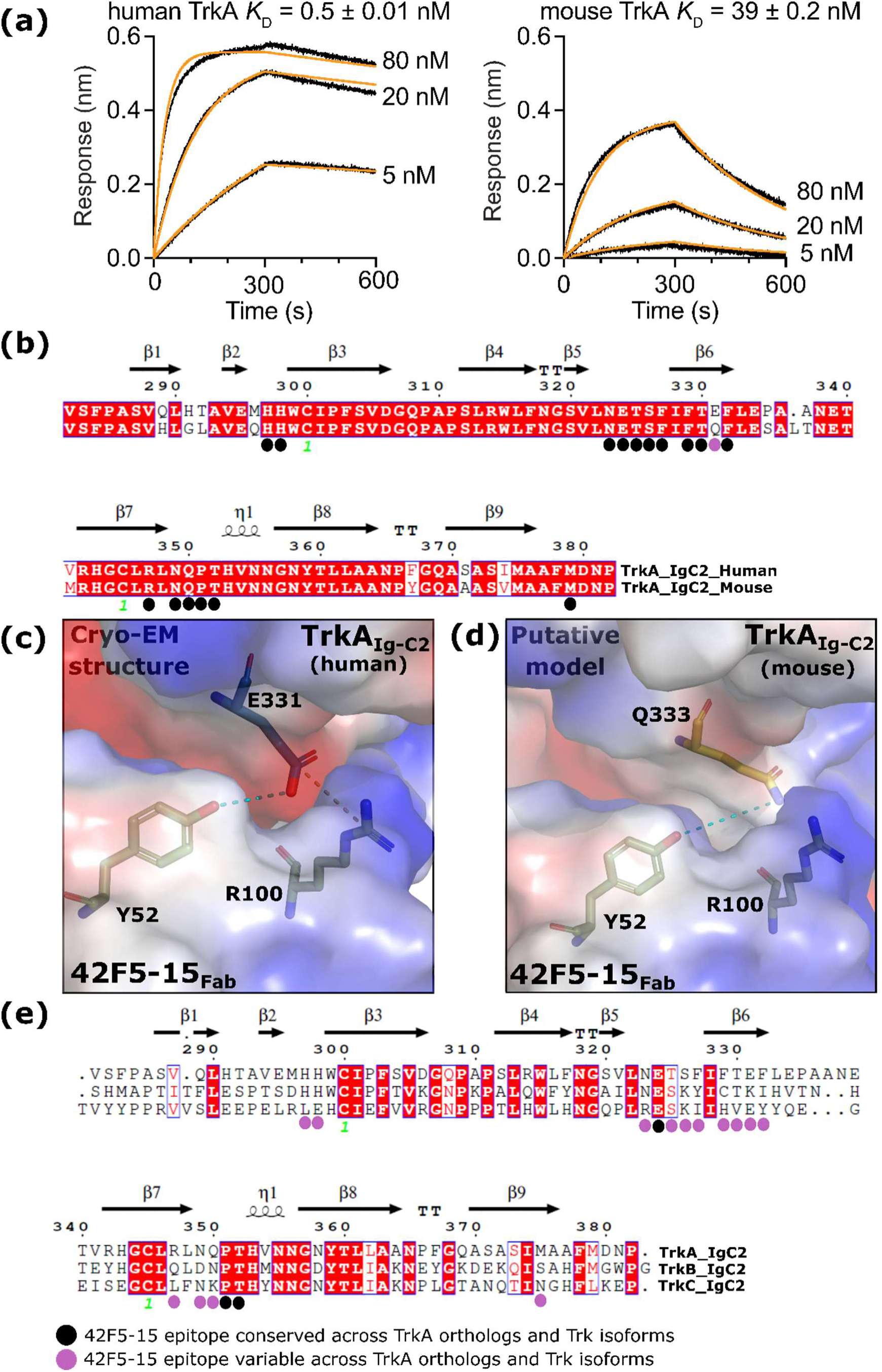
Basis for species-dependent TrkA affinity and structural determinants of Trk isoform selectivity of the mAb 42F5-15. **(a)** Binding affinities of mAb 42F5-15 measured for human and mouse TrkA reveals strong affinity for human ortholog. **(b)** Structure based sequence alignment of TrkA orthologs (human and mouse) generated by superposition of structures of TrkA IgC2 domain is shown. Qualitative electrostatic surface map of TrkA Ig-C2 and 43F5-15 Fab is drawn to illustrate the presence of charge complementarity between **(c)** E331-R100 pair at TrkA_ECD_/42F5-15_Fab_ interface (human) and absence of the same between **(d)** Q333-R100 pair at putative TrkA_ECD_/42F5-15_Fab_ interface (mouse). The red and blue color of electrostatic surface map corresponds to negative and positive charges, respectively. Dashed lines at the at the TrkA_ECD_/42F5-15_Fab_ interface indicate hydrogen bonds (cyan) and ionic (orange) interactions. **(e)** Structure based sequence alignment of Trk isoforms (TrkA, TrkB, and TrkC) generated by superposition of structures of Trk IgC2 domain is shown. **(b, e)** Conserved residues are marked in white on a red background while similar residues are in red. Residues in blue frame depict similarity across groups. Secondary structural elements and residue numbering indicated above the alignment are extracted from the structure of TrkA IgC2 domain as observed in TrkA_ECD_/42F5-15_Fab_ complex. TrkA IgC2 residues that bind 42F5-15_Fab_ (epitope), as inferred from structural analysis of TrkA_ECD_/42F5-15_Fab_ complex, are indicated by colored circles below the alignment. Black and pink circles indicate conserved and variable residues at mAb 42F5-15 epitope positions across TrkA orthologs **(b)** and Trk isoforms **(e).**

Analysis of the cryo-EM structure of the human TrkA_ECD_/42F5-15_Fab_ complex shows that residue E331 forms a complex, bidentate interaction network at the interface, consisting of a hydrogen bond with Y52 of CDR H2 and a salt bridge with R100 of CDR H3 on the mAb 42F5-15 (**Fig. 2b, 4c**). In mouse TrkA, E331 is substituted by a Glutamine (Q333). Given that Glutamine is nearly isosteric to Glutamate but lacks a formal negative charge, it is tempting to hypothesize that in the putative mouse TrkA_ECD_/42F5-15_Fab_ complex, Q333 of mouse TrkA may retain the hydrogen bond with Y52 of CDR H2 but suffers a total loss of the interfacial electrostatic salt bridge with the R100 of CDR H3 (**Fig. 4d**).

mAb 42F5-15 binds TrkA but not TrkB or TrkC [33]. The epitope of 42F5-15 on TrkA is not conserved across TrkB and TrkC, with multiple non-isosteric substitutions (**Fig. 4e**). Notably, if a single isosteric glutamate-to-glutamine substitution between human and mouse TrkA (**Fig. 4b**) results in a ∼ 80-fold reduction in the mAb binding affinity to mouse TrkA, then the presence of multiple non-isosteric substitutions across the corresponding epitope positions of TrkB and TrkC (**Fig. 4e**) would be expected to impose even greater disruption of electrostatic and shape complementarity at the antibody-receptor interface. This provides a structural basis for the high selectivity of the mAb for the TrkA isoform, mediating disruption of NGF/TrkA interaction without perturbing TrkB and TrkC signaling pathways [33].

### 6. mAb 42F5-15 abrogates NGF-induced mechanical allodynia in mice

NGF is known to induce pain in both humans and rodents [5]. The mAb 42F5-15 inhibits TrkA-mediated signaling and increases pain threshold [33]. To independently confirm that the TrkA antibody treatment counteracts NGF-induced nociception, male and female mice were administered TrkA antibody or control IgG1 intraperitoneally (i.p., 1.67 mg/kg). 30 minutes later, mice received an intraplantar (i.pl.), injection of NGF or vehicle, (50 ng/10 µL), and mechanical allodynia was assessed beginning 30 minutes after the second injection (**Fig. 5a**).

**Figure 5.**
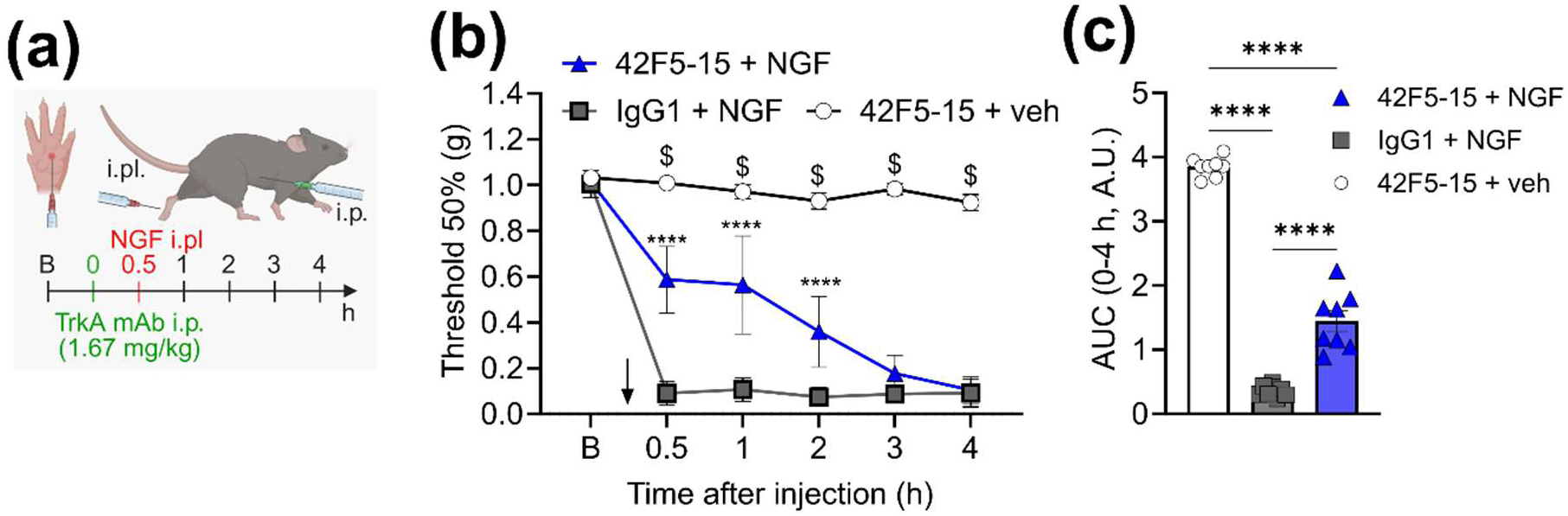
TrkA antibody abrogates NGF-induced mechanical allodynia in mice. **(a)** Schematic representation of TrkA antibody and NGF injections, and mechanical allodynia assessment. **(b)** Effect of TrkA antibody (1.67 mg/kg, i.p.) in male and female mice. After baseline (B) measurements, the TrkA antibody or IgG1 control was injected. 30 min later, mouse NGF (50 ng/10 μl, i.pl.) was administered. Mechanical allodynia was measured. n=8. **(c)** Area under curve (AUC) of time courses. Mean±SEM. \*\*\*\**P*<0.0001 vs. IgG1, $*P*<0.0001 vs. Vehicle (Veh) **(b)** 2-way ANOVA, Tukey’s multiple comparisons. **(c)** 1-way ANOVA, Tukey’s multiple comparisons.

Systemic administration of TrkA antibody substantially reduced NGF-induced mechanical allodynia up to 2 h after intraplantar injection. NGF injection, compared with IgG1 control-treated mice (**Fig. 5b, c**). The nociceptive effect of NGF was local, as no significant change in withdrawal threshold was observed in the contralateral paw of NGF-injected mice (**Fig. S1**). In vehicle-injected animals, TrkA antibody did not alter mechanical withdrawal thresholds in either the ipsilateral or contralateral paw (**Fig. 5B, Fig. S1**). Together, these results indicate that this anti-TrkA antibody efficiently counteracts NGF-induced nociceptive sensitization, supporting its efficacy in reducing NGF-induced mechanical allodynia.

**Supplementary Fig. S1.**
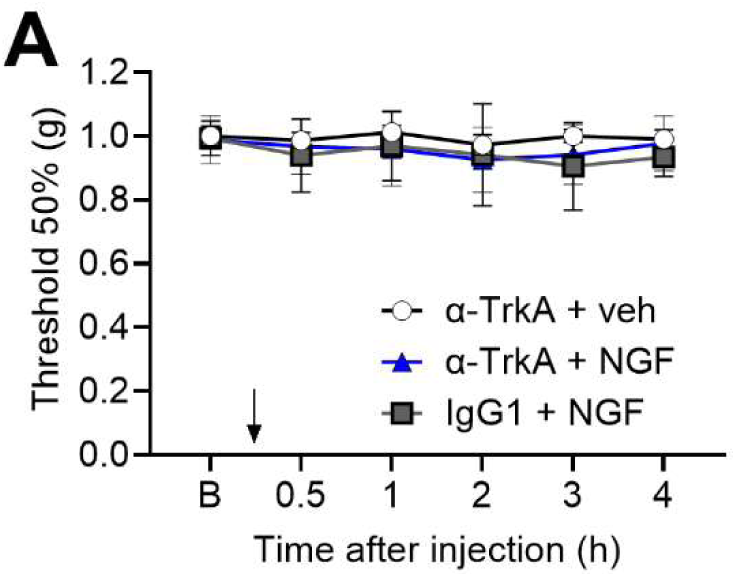
Withdrawal responses of the contralateral paw. Withdrawal responses of the contralateral paw to VFF stimulation after injection of NGF into the ipsilateral hindpaw. N=8. 2-way ANOVA, Tukey’s multiple comparisons.

## Discussion

TrkA, the high-affinity receptor for nerve growth factor (NGF), is a central regulator of neuronal survival during embryogenesis and nociceptive signaling in the adult peripheral nervous system [1,17]. The NGF/TrkA signaling axis therefore represents a key pathway in the modulation of pain and has emerged as a clinically validated therapeutic target for the development of novel analgesics [14,15]. However, the development of anti-NGF biologics has been limited by safety concerns associated with global ligand sequestration [22]. Consequently, the therapeutic landscape for targeting this pathway has shifted toward receptor-specific strategies [32] aimed at selectively disrupting NGF/TrkA interactions while preserving NGF availability. Targeting the extracellular domain of the TrkA receptor with monoclonal antibodies offers a more localized and selective approach. Unlike anti-NGF antibodies, anti-TrkA antibodies do not sequester NGF, potentially allowing for more nuanced modulation of the pathway. Elucidating the structural basis of this TrkA ECD-mediated inhibition is therefore essential for defining how receptor-directed antagonists modulate the NGF/TrkA signaling axis. While MNAC13 is a well-characterized neutralizing anti-TrkA antibody that has been shown to reduce inflammatory and neuropathic pain in several mouse models, its exact binding site on TrkA and neutralization mechanism remains poorly understood due to the absence of a TrkA_ECD_/MNAC13_Fab_ complex structure [31,32].

Recently, use of TrkA neutralizing antibody 42F5-15 has been proposed as a strategy for developing drugs to treat pain including inflammatory, neuropathic, cancer and osteoarthritis [33]. In the present manuscript, we report the high-resolution structure of anti-TrkA mAb 42F5-15 in complex with the TrkA extracellular domain (ECD). The structural analysis revealed that the mAb 42F5-15 neutralizes NGF/TrkA signaling by acting as a direct competitive antagonist of NGF. By physically occupying critical NGF binding hotspot on TrkA, mAb 42F5-15 precludes the initial binding of NGF to the receptor, providing a robust and unambiguous mechanism for neutralization that ensures that TrkA remains inaccessible to NGF, thus preventing formation of NGF/TrkA complex. Since, formation of NGF/TrkA complex is essential for downstream signaling known to orchestrate pain [15], the mAb 42F5-15 provides *in vivo* neutralizing efficacy against the NGF/TrkA axis in pain-related behavior [33]. In contrast, the AlphaFold3 [35] prediction of TrkA_ECD_/MNAC13_Fab_ complex, supported by previously reported epitope mapping studies [32], indicates that MNAC13 binds to TrkA IgC1 domain and is not structurally positioned to interfere with the initial docking of NGF on TrkA. This observation aligns with previous competition experiments between MNAC13 and NGF for TrkA ECD binding showed that *in vitro* binding of MNAC13 to the soluble TrkA ECD is not inhibited by NGF and vice versa [32]. Its *in vivo* anti-TrkA activity is hypothesized to be allosteric mediation [32]. The activity of the allosterically neutralizing mAb MNAC13 will depend on modulation of specific conformational transitions or receptor states, which may vary across cellular and physiological contexts. In contrast to allosteric inhibitors, whose efficacy may depend on receptor conformational dynamics and ligand concentration, orthosteric antagonist antibodies directly compete for the primary ligand-binding interface and sterically prevent ligand engagement. This suggests that the mAb 42F5-15 offer superior neutralizing potency [33], particularly in high-ligand environments where allosteric inhibitors like MNAC13 might be bypassed. Targeting the primary functional NGF/TrkA interface rather than an accessory regulatory region may represent a mechanistically advantageous strategy for achieving robust inhibition of NGF/TrkA signaling [33].

While small-molecule inhibitors targeting the intracellular TrkA kinase domain have been extensively developed [26,27], primarily for oncological applications, their therapeutic utility in chronic pain management remains limited by the challenge of achieving high receptor isoform selectivity over TrkB and TrkC, which possess highly conserved ATP-binding pockets [8]. Because TrkB and TrkC mediated signaling pathways are essential for central nervous system homeostasis, particularly in cognition and motor function, off-target kinase inhibition can lead to neurological adverse effects [28]. mAb 42F5-15 is shown to bind TrkA but not TrkB or TrkC [33]. Our comparative structural analysis demonstrates that the mAb 42F5-15 targets epitope composed of residues uniquely conserved in TrkA IgC2 domain but structurally divergent in TrkB and TrkC in IgC2 domains, thereby defining the molecular determinants underlying its isoform selectivity [33].

The high-resolution cryo-EM structure enabled epitope mapping and Fab modeling at near-atomic resolution, identifying key molecular interactions and CDR residues contributing to binding affinity and energetics. The structure provides a framework to guide antibody engineering for improved pharmacokinetic properties while preserving potency [36]. While biologics such as Fabs offer high specificity, small molecules have better oral bioavailability. High-resolution structural mapping of the TrkA_ECD_/42F5-15_Fab_ interface provides the necessary framework for virtual screening and fragment-based drug discovery, enabling identification of small molecules that mimic the inhibitory effect of the mAb 42F5-15.

*NTRK1* gene fusions represent oncogenic drivers across a range of malignancies including lung adenocarcinoma, thyroid carcinoma, and pediatric gliomas [37]. These fusion proteins are predominantly intracellular signaling kinases activated through partner-mediated dimerization with current therapeutic approaches relying primarily on small-molecule kinase inhibitors [26,27]. While such kinase-targeted therapies have demonstrated clinical efficacy, resistance mutations and on-target toxicities highlight the need for alternative or complementary strategies. In this context, the high-resolution structural characterization of the TrkA ECD in complex with the neutralizing antibody provides a framework for targeting full-length, cell-surface TrkA ECD in tumor settings where wild-type receptor expression is retained or upregulated [38], including tumor subpopulations, stromal compartments, or tumor-nerve interface niches where NGF/TrkA signaling may contribute to tumor biology or pain. Importantly, although *NTRK1* fusion proteins themselves are not accessible to extracellular antibodies, TrkA ECD-directed antibodies can be leveraged for antibody-drug conjugate (ADC) development [39] in TrkA-positive malignancies expressing surface full-length receptor [38], enabling selective delivery of cytotoxic payloads to tumor cells with validated TrkA surface expression. Furthermore, structural insight from the TrkA_ECD_/42F5-15_Fab_ interface enables rational engineering of antibody scaffolds with improved affinity, internalization properties, and epitope specificity, which are key determinants of antibody-drug conjugate efficacy. In addition, such antibodies may provide a platform for combination strategies with Trk kinase inhibitors, potentially addressing both extracellular signaling and intracellular oncogenic driver activity within heterogeneous tumor ecosystems.

## Conclusion

In the absence of other TrkA-mAb complex structures in the PDB, this work provides structural characterization of a high-affinity monoclonal antibody against the TrkA receptor. By elucidating the molecular basis for disruption of the NGF/TrkA interaction, the antibody is shown to prevent the formation of the NGF/TrkA complex, a critical initial step in the downstream pain-signaling cascades, by direct steric blockade of the NGF binding to TrkA Ig-C2 domain. By providing a high-resolution view of selective inhibition of NGF/TrkA complex formation by the anti-TrkA mAb, this structure reveals a mechanistic basis for receptor-directed antagonism that enables more selective pain modulation compared with systemic NGF sequestration [33].

Collectively, the high resolution cryo-EM structure of TrkA_ECD_/42F5-15_Fab_ complex provides a structural framework for exploiting TrkA extracellular domain recognition as a platform for selective modulation of TrkA biology across disease contexts. In the setting of pain, this enables receptor-directed intervention within the NGF/TrkA signaling axis to attenuate nociceptive sensitization without global ligand sequestration [33]. In oncology, this approach may be leveraged to target tumor compartments and tumor-microenvironment interfaces that retain surface TrkA expression, supporting antibody-based therapeutic modalities that can complement intracellular kinase inhibition strategies in TrkA-driven malignancies.

## Methods

### TrkA DNA construct

TrkA construct contains a N-terminal glycoprotein 64 (gp64) signal peptide for extracellular secretion followed by the TrkA-ECD (extracellular domain from amino acid residues 33-415 (UniProt #P04629-1)) and a C-terminal 10x His-tag in a pFastBac vector (ThermoFisher).

### Expression and purification of TrkA

Transfer vectors containing TrkA was transformed into chemically competent *E.coli* EmBacY cells (Geneva Biotech, Switzerland) and recombinant baculovirus DNA was extracted by conventional miniprep method. Five mg of DNA was transfected into Sf9 insect cells and supernatant containing the “P1” virus was collected after five days. P1 virus was used to infect Sf9 cells to generate “P2” virus stock.

For expression of TrkA, 300-600 ml of Sf9 cells at 1×10^6^ cells/ml were infected with 3-6 ml of P2 stock (1:100 dilution) and cells were allowed to grow at 27°C for 5-7 days. Clarified media was then loaded onto PROTEINDEX™ Ni-Penta™ columns (Marvelgent Biosciences, Waltham, MA), washed and proteins were eluted with an imidazole gradient from 5-500 mM. Fractions containing TrkA protein, as visualized by Coomassie-stained protein gels, were pooled, concentrated and further purified by size-exclusion chromatography using a Superdex 200 Increase 10/300 GL (Cytiva) in 40 mM HEPES, pH=7.8; 200 mM NaCl If necessary, fractions containing protein were concentrated to a concentration of at least 0.5 mg/ml. Aliquoted samples were flash-frozen and stored at −80°C.

### Cryo-EM sample preparation and data collection

The amino acid sequence of the V_L_ and V_H_ domains of anti-TrkA antibody 42F5-15 is from a published patent application [33]. The full IgG sequence was designed using the human IgG1 sequence and the Fc-silencing LALA-PG mutations [40]. Antibody production was performed by Genscript (Piscataway, NJ). The Fab sample was generated using the Thermo Scientific Pierce Fab Micro Preparation Kit (Thermo Fisher Scientific) according to the manufacturer’s protocol. The kit uses immobilized papain protease to digest human or mouse IgG antibodies to make separate Fab and Fc regions and subsequently to purify the Fab using Protein A agarose. The purified Fab was subsequently incubated with TrkA ectodomain (ECD) at a 1.3:1 molar ratio at 4°C for ∼ 1 hour to assemble the TrkA_ECD_/42F5-15_Fab_ complex. 3 μL of that solution, at the concentration of ∼0.6 mg/ml, was applied to the plasma cleaned UltraAuFoil holey gold grids (Quantifoil R 1.2/1.3). Using a Vitrobot Mark IV (Thermo Fisher), the grids were blotted with the following settings: 4°C, 100% humidity, 2.0 sec blot time, blot force 10 and plunge frozen in liquid ethane pre-cooled by liquid nitrogen [41]. Micrographs were recorded using an EF-Krios (Thermo Fisher) electron microscope operating at 300 kV and equipped with a Gatan K3 camera with a nominal magnification of 105,000X and calibrated with the pixel size of 0.4125 Å. Movies were collected using Leginon v 3.7 [42] at a dose rate of 25.43 e-/Å^2^/s with a total exposure of 1.80 seconds, for an accumulated dose of 45.77 e-/Å^2^. Intermediate frames were recorded every 0.05 seconds for a total of 40 frames per micrograph. A total of 8086 movies were collected at a nominal defocus range of 0.6 – 2.5 μm. Ice thickness was determined as described in [43].

### Cryo-EM data processing

#### Particle picking

Motion correction and CTF estimation of the collected movies were done in CryoSPARC [44] using Patch Motion Correction and Patch CTF Estimation, respectively. Since the movies were collected in super-resolution mode, the output F-crop factor was set to ½ in Patch Motion Correction. Micrographs were then curated by selecting only those with CTF Fit Resolution < 4 Å and Total Full Frame Motion < 200 pixels in Manually Curate Exposures job. The manually curated micrographs were then subjected to Micrograph Junk Detector and Micrograph Denoiser jobs in CryoSPARC to minimize picking of junk particles and enhance picking of TrkA_ECD_/42F5-15_Fab_ complex particles. Initially, Blob Picker was used to pick particles of TrkA_ECD_/42F5-15_Fab_ complex from the 1000 denoised micrographs. Since the stoichiometry and the diameter of the TrkA_ECD_/42F5-15_Fab_ complex was not known, a particle diameter range of 120 Å, was used in Blob Picker. Micrograph Junk Detector was used to reject particles picked from and around junk regions. Inspect Particle Picks job was then used in Auto Cluster mode to remove false positive particle picks and select the cluster of candidate particles. The resulting good particles were then extracted at a box size of 360 pixels. 2D Classification job was run with 200 classes, 160 Å circular mask diameter, 40 number of online-EM iterations, 400 batchsize per class. Select 2D was used to select classes consistent with TrkA_ECD_ domain architecture. The selected 2D classes were used as templates to pick particles from all the denoised micrographs using Template Picker. The template picked particles were again passed through Micrograph Junk Detector and Inspect Particle Picks to filter junk particles and false positives. The resulting good particles were then extracted at a box size of 360 pixels and carried downstream to particle curation.

#### Particle curation

The initial 2D Classification results of the blob picked particles revealed 2D class averages resembling Fab/TrkA_ECD_ complex. Since there were no available 3D references, the particle curation of the template picked particles was first done in 2D. 2D Classification job on template picked particles was run with 250 classes, 160 Å circular mask diameter, 80 number of online-EM iterations, 400 batchsize per class. 2D Classification results revealed 2D class averages with 1:1 TrkA_ECD_/42F5-15_Fab_. Ab-initio Reconstruction jobs were run to generate candidate/representative 3D references from the sets of 2D class averages. The ab-initio generated 3D references, with apparent topology resembling Fab/TrkA_ECD_ complex, were then used for particle curation in 3D. An initial round of particle cleanup was done using separate heterogenous refinement jobs with ab-initio selected and decoy 3D maps as input volumes. These particles which did not get sorted to decoy maps were further 3D classified using heterogenous refinement. Each of the classes from the heterogenous refinement was then refined further using Non-Uniform refinement [45] job to improve the resolution. The 3D classification and refinement yielded distinct 3D reconstructions at resolutions 2.90 Å, 2.70 Å, and 2.60 Å, based on gold-standard Fourier shell correlation of 0.143 criterion, that together provide a multi-scale view of the TrkA_ECD_/42F5-15_Fab_ complex **(Fig. 1).** Focused refinements with masks corresponding to the TrkA_ECD_ or 42F5-15_Fab_ or TrkA_ECD_/42F5-15_Fab_ were done to improve the quality or the resolution of the maps. The 2.90 Å and 2.60 Å maps were utilized for atomic model building and analysis of TrkA_ECD_/42F5-15_Fab_ complex.

### Cryo-EM structure modeling, refinement and analysis

For modeling TrkA_ECD_ and 42F5-15_Fab_ in TrkA_ECD_/42F5-15_Fab_ density maps, initial coordinates for TrkA_ECD_ were taken from TrkA/NGF crystal structure (PDB 2IFG) while 42F5-15_Fab_ sequences were used to predict an AlphaFold3 [35] model of 42F5-15_Fab_ to provide starting set of coordinates. Initial coordinates were first rigid-body fitted into the cryo-EM density maps using UCSF ChimeraX [46]. The rigid body fitting of TrkA_ECD_ and 42F5-15_Fab_ in TrkA_ECD_/42F5-15_Fab_ density maps in cryo-EM density maps was followed by iterative rounds of real space refinement and manual model building in PHENIX [47], COOT [48] and ISOLDE [49]. The validated and refined models were used for molecular analysis of TrkA_ECD_/42F5-15_Fab_ complex and figure generation was done using UCSF ChimeraX [46]. Surface area calculations and the identification of interface residues were performed using PDBePISA [50]. PLIP standalone tool [51] was used for calculations of molecular interactions at the interface of the TrkA_ECD_/42F5-15_Fab_ complex.

### Biolayer interferometry (BLI)

BLI measurements were performed on an Octet RED96e (Sartorius), at 30°C in 50 mM Tris HCl buffer, pH 7.5, containing 150 mM NaCl, 0.005% Tween-20, 0.5% BSA, and 2 µM biotin. The anti-TrkA antibody was immobilized on AHC2 biosensors (Sartorius) to a level of approximately 0.5 nm. Mouse TrkA Protein was purchased from MedChemExpress (#HY-P76116). Data were double-referenced and fit using a global 1:1 binding model in Octet Data Analysis HT v.12.0.2.59.

### Animals

Experiments were in accordance with the guidelines recommended by the National Institute of Health, the International Association for the study of Pain, the National Centre for the Replacement, Refinement and Reduction of Animals in Research (ARRIVE) guidelines, and were approved by the New York University Institutional Animal Care and Use Committee (PROTO202000006). Male and female C57BL/6 mice (8-10 weeks for behavioral experiments, Jackson Laboratory) were used. Animals were housed five per cage at 22 ± 0.5°C under a controlled 14/10 h light/dark cycle with free access to food and water. Mice were randomly assigned to experimental groups; group size was based on previous similar studies. Investigators were blind to treatments.

### Intraplantar NGF injection model of acute pain

Mice were anesthetized (isofluorane) and received intra-plantrar injection of NGF (50 ng/10 µL; cat# 1156-NG-100/CF, R&D) or vehicle (0.9% NaCl, 10 µL) as a control [17]. Animals were habituated to the pain behavioral tests before baselines were taken. Nociceptive behavior was assessed in the morning prior to NGF or vehicle injection. Nociception assays were assessed 30 minutes post model induction.

### Mechanical allodynia assessment

Mechanical allodynia was evaluated by measuring hindpaw withdrawal response to von Frey filaments stimulation using the up-and-down method [52]. The 50% threshold was determined by the following equation: 50% threshold (g) = 10 log (last filament) + k × δ. The constant, k, was found in the table by Dixon [53] and determined by the response pattern.

### Statistical analysis

Statistical analyses were performed using GraphPad Prism (v11.0.1). Data distribution was assessed using the D’Agostino–Pearson normality test. Data are shown as mean ± SEM. Statistical significance was set at *P*<0.05. For comparisons involving more than two groups, 1-way ANOVA followed by Tukey’s post hoc test for normally distributed data. 2-way ANOVA followed by Tukey’s post hoc tests was used for multi-factorial analyses as appropriate.

### Data availability

Cryo-EM maps and models generated in this study for TrkA_ECD_/42F5-15_Fab_ complex have been deposited in the EMDB and PDB database respectively. Statistics regarding cryo-EM data collection, processing, structure refinement and validation are given in Table 1.

**Table 1.**
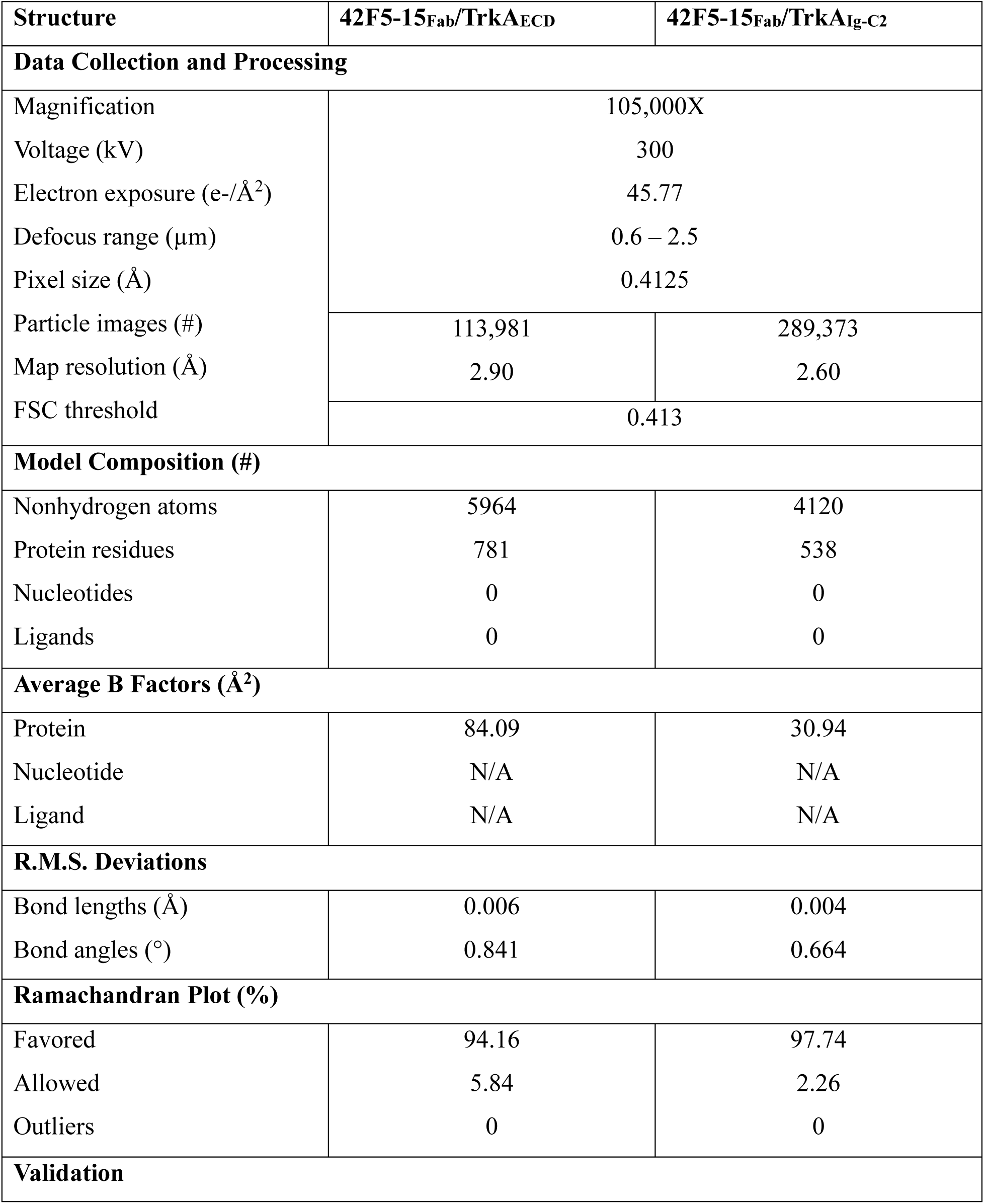
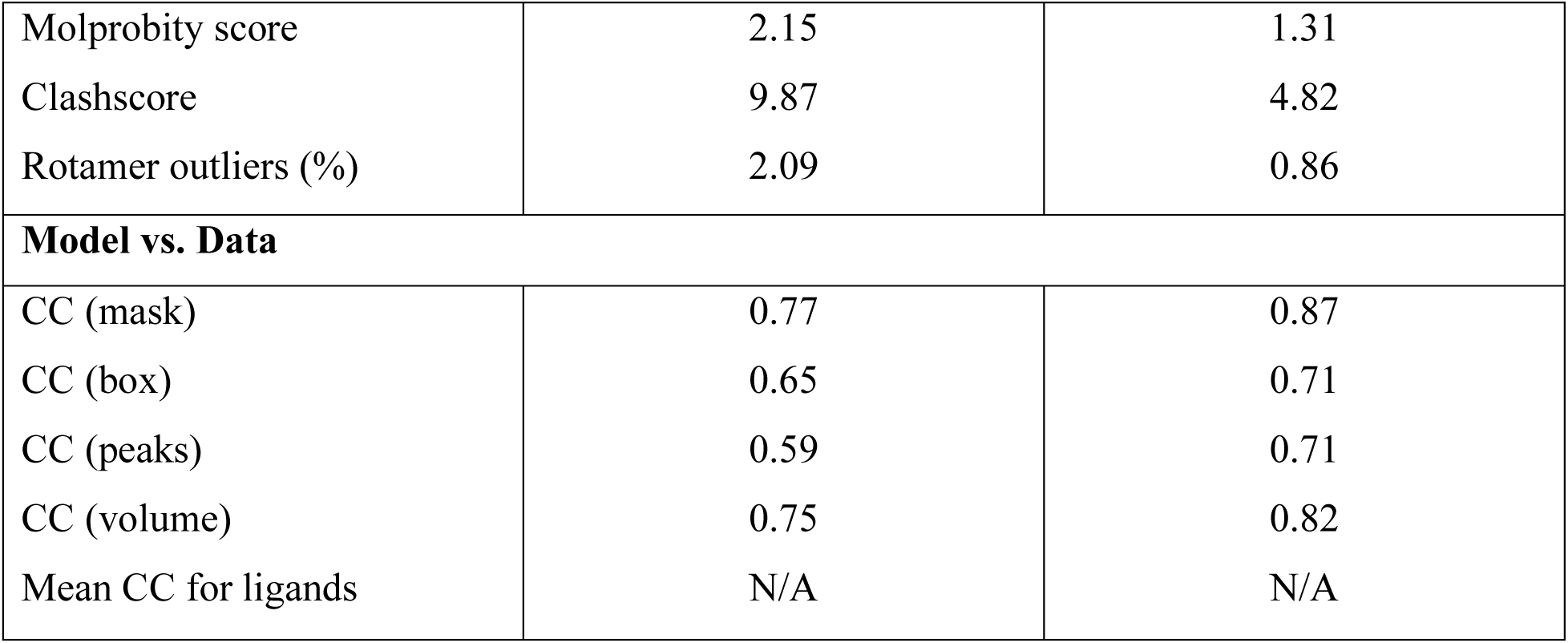
Parameters and statistics for cryo-EM data collection, processing, structure refinement, and validation.

## References

1. Fagan, A.M. et al. (1996) TrkA, but not TrkC, receptors are essential for survival of sympathetic neurons in vivo. J Neurosci 16, 6208–6218. 10.1523/jneurosci.16-19-06208.1996

2. Ehlers, M.D. et al. (1995) NGF-stimulated retrograde transport of trkA in the mammalian nervous system. J Cell Biol 130, 149–156. 10.1083/jcb.130.1.149

3. Smeyne, R.J. et al. (1994) Severe sensory and sympathetic neuropathies in mice carrying a disrupted Trk/NGF receptor gene. Nature 368, 246–249. 10.1038/368246a0

4. Barker, P.A. et al. (2020) Nerve Growth Factor Signaling and Its Contribution to Pain. J Pain Res 13, 1223–1241. 10.2147/jpr.S247472

5. Denk, F. et al. (2017) Nerve Growth Factor and Pain Mechanisms. Annu Rev Neurosci 40, 307–325. 10.1146/annurev-neuro-072116-031121

6. Woolf, C.J. et al. (1994) Nerve growth factor contributes to the generation of inflammatory sensory hypersensitivity. Neuroscience 62, 327–331. 10.1016/0306-4522(94)90366-2

7. Wehrman, T. et al. (2007) Structural and Mechanistic Insights into Nerve Growth Factor Interactions with the TrkA and p75 Receptors. Neuron 53, 25–38. 10.1016/j.neuron.2006.09.034

8. Huang, E.J. and Reichardt, L.F. (2003) Trk Receptors: Roles in Neuronal Signal Transduction*. Annual Review of Biochemistry 72, 609–642. 10.1146/annurev.biochem.72.121801.161629

9. Zhang, X. et al. (2005) NGF rapidly increases membrane expression of TRPV1 heat-gated ion channels. Embo j 24, 4211–4223. 10.1038/sj.emboj.7600893

10. Ji, R.-R. et al. (2002) p38 MAPK Activation by NGF in Primary Sensory Neurons after Inflammation Increases TRPV1 Levels and Maintains Heat Hyperalgesia. Neuron 36, 57–68. 10.1016/S0896-6273(02)00908-X

11. Marlin, M.C. and Li, G. (2015) Biogenesis and function of the NGF/TrkA signaling endosome. Int Rev Cell Mol Biol 314, 239–257. 10.1016/bs.ircmb.2014.10.002

12. Donnerer, J. et al. (1992) Increased content and transport of substance P and calcitonin gene-related peptide in sensory nerves innervating inflamed tissue: evidence for a regulatory function of nerve growth factor in vivo. Neuroscience 49, 693–698. 10.1016/0306-4522(92)90237-v

13. Gould, H.J., 3rd et al. (2000) A possible role for nerve growth factor in the augmentation of sodium channels in models of chronic pain. Brain Res 854, 19–29. 10.1016/s0006-8993(99)02216-7

14. Mantyh, P.W. et al. (2011) Antagonism of nerve growth factor-TrkA signaling and the relief of pain. Anesthesiology 115, 189–204. 10.1097/ALN.0b013e31821b1ac5

15. Hirose, M. et al. (2016) NGF/TrkA Signaling as a Therapeutic Target for Pain. Pain Pract 16, 175–182. 10.1111/papr.12342

16. Mardy, S. et al. (1999) Congenital Insensitivity to Pain with Anhidrosis: Novel Mutations in the TRKA (NTRK1) Gene Encoding A High-Affinity Receptor for Nerve Growth Factor. The American Journal of Human Genetics 64, 1570–1579. 10.1086/302422

17. Peach, C.J. et al. (2024) Neuropilin-1 inhibition suppresses nerve growth factor signaling and nociception in pain models. J Clin Invest 135. 10.1172/jci183873

18. Wangzhou, A. and Price, T.J. (2025) Reinvigorating drug development around NGF signaling for pain. J Clin Invest 135. 10.1172/jci189029

19. Jayabalan, P. and Schnitzer, T.J. (2017) Tanezumab in the treatment of chronic musculoskeletal conditions. Expert Opin Biol Ther 17, 245–254. 10.1080/14712598.2017.1271873

20. DiMartino, S.J. et al. (2025) Efficacy and safety of fasinumab in an NSAID-controlled study in patients with pain due to osteoarthritis of the knee or hip. BMC Musculoskelet Disord 26, 192. 10.1186/s12891-025-08402-8

21. Lane Nancy, E. et al. Tanezumab for the Treatment of Pain from Osteoarthritis of the Knee. New England Journal of Medicine 363, 1521–1531. 10.1056/NEJMoa0901510

22. Hochberg, M.C. (2015) Serious joint-related adverse events in randomized controlled trials of anti-nerve growth factor monoclonal antibodies. Osteoarthritis and Cartilage 23, S18–S21. 10.1016/j.joca.2014.10.005

23. Baeza-Raja, B. et al. (2016) p75 Neurotrophin Receptor Regulates Energy Balance in Obesity. Cell Reports 14, 255–268. 10.1016/j.celrep.2015.12.028

24. Toni, T. et al. (2014) Systems Pharmacology of the NGF Signaling Through p75 and TrkA Receptors. CPT Pharmacometrics Syst Pharmacol 3, e150. 10.1038/psp.2014.48

25. Xu, J. et al. NGF-p75 signaling coordinates skeletal cell migration during bone repair. Science Advances 8, eabl5716. 10.1126/sciadv.abl5716

26. Liu, D. et al. (2018) Entrectinib: an orally available, selective tyrosine kinase inhibitor for the treatment of NTRK, ROS1, and ALK fusion-positive solid tumors. Ther Clin Risk Manag 14, 1247–1252. 10.2147/tcrm.S147381

27. Federman, N. and McDermott, R. (2019) Larotrectinib, a highly selective tropomyosin receptor kinase (TRK) inhibitor for the treatment of TRK fusion cancer. Expert Rev Clin Pharmacol 12, 931–939. 10.1080/17512433.2019.1661775

28. Giustini, N.P. et al. (2022) Development of Neuropathic Arthropathy With Entrectinib: Case Report. JTO Clinical and Research Reports 3. 10.1016/j.jtocrr.2022.100419

29. Urfer, R. et al. (1998) High Resolution Mapping of the Binding Site of TrkA for Nerve Growth Factor and TrkC for Neurotrophin-3 on the Second Immunoglobulin-like Domain of the Trk Receptors*. Journal of Biological Chemistry 273, 5829–5840. 10.1074/jbc.273.10.5829

30. Wiesmann, C. et al. (1999) Crystal structure of nerve growth factor in complex with the ligand-binding domain of the TrkA receptor. Nature 401, 184–188. 10.1038/43705

31. Ugolini, G. et al. (2007) The function neutralizing anti-TrkA antibody MNAC13 reduces inflammatory and neuropathic pain. Proceedings of the National Academy of Sciences 104, 2985–2990. 10.1073/pnas.0611253104

32. Covaceuszach, S. et al. (2005) Neutralization of NGF-TrkA receptor interaction by the novel antagonistic anti-TrkA monoclonal antibody MNAC13: A structural insight. Proteins: Structure, Function, and Bioinformatics 58, 717–727. 10.1002/prot.20366

33. Yuan, X. et al. (2026). Anti-trka antibody or antigen-binding fragment thereof, preparation method thereof, and application thereof. U.S. Patent No. 12,590,158.

34. Ramaraj, T. et al. (2012) Antigen–antibody interface properties: Composition, residue interactions, and features of 53 non-redundant structures. Biochimica et Biophysica Acta (BBA) - Proteins and Proteomics 1824, 520–532. 10.1016/j.bbapap.2011.12.007

35. Abramson, J. et al. (2024) Accurate structure prediction of biomolecular interactions with AlphaFold 3. Nature 630, 493–500. 10.1038/s41586-024-07487-w

36. Riso, M. et al. (2025) Binding mode–guided development of high-performance antibodies targeting site-specific posttranslational modifications. Proceedings of the National Academy of Sciences 122, e2411720121. 10.1073/pnas.2411720121

37. Joshi, S.K. et al. (2019) Revisiting NTRKs as an emerging oncogene in hematological malignancies. Leukemia 33, 2563–2574 10.1038/s41375-019-0576-8

38. Jagana, H.L. et al. (2026) Oncogenic influences of neurotrophin receptors: Shedding light on Trk biology. Cell Reports 45. 10.1016/j.celrep.2026.116928

39. Hattori, T. et al. (2025) Engineering antibody-drug conjugates targeting an adhesion GPCR, CD97. Proc Natl Acad Sci U S A 122, e2516627122. 10.1073/pnas.2516627122

40. Lo, M. et al. (2017) Effector-attenuating Substitutions That Maintain Antibody Stability and Reduce Toxicity in Mice. J Biol Chem 292, 3900–3908. 10.1074/jbc.M116.767749

41. Dubochet, J. et al. (1988) Cryo-electron microscopy of vitrified specimens. Q Rev Biophys 21, 129–228 10.1017/s0033583500004297

42. Cheng, A. et al. (2021) Leginon: New features and applications. Protein Science 30, 136–150. 10.1002/pro.3967

43. Rice, W.J. et al. (2018) Routine determination of ice thickness for cryo-EM grids. J Struct Biol 204, 38–44. 10.1016/j.jsb.2018.06.007

44. Punjani, A. et al. (2017) cryoSPARC: algorithms for rapid unsupervised cryo-EM structure determination. Nature Methods 14, 290–296. 10.1038/nmeth.4169

45. Punjani, A. et al. (2020) Non-uniform refinement: adaptive regularization improves single-particle cryo-EM reconstruction. Nature Methods 17, 1214–1221. 10.1038/s41592-020-00990-8

46. Pettersen, E.F. et al. (2021) UCSF ChimeraX: Structure visualization for researchers, educators, and developers. Protein Sci 30, 70–82. 10.1002/pro.3943

47. Afonine, P.V. et al. (2018) Real-space refinement in PHENIX for cryo-EM and crystallography. Acta Crystallogr D Struct Biol 74, 531–544. 10.1107/s2059798318006551

48. Emsley, P. and Cowtan, K. (2004) Coot: model-building tools for molecular graphics. Acta Crystallogr D Biol Crystallogr 60, 2126–2132 10.1107/s0907444904019158

49. Croll, T.I. (2018) ISOLDE: a physically realistic environment for model building into low-resolution electron-density maps. Acta Crystallogr D Struct Biol 74, 519–530. 10.1107/s2059798318002425

50. Krissinel, E. and Henrick, K. (2007) Inference of macromolecular assemblies from crystalline state. J Mol Biol 372, 774–797. 10.1016/j.jmb.2007.05.022

51. Schake, P. et al. (2025) PLIP 2025: introducing protein–protein interactions to the protein–ligand interaction profiler. Nucleic Acids Research 53, W463–W465. 10.1093/nar/gkaf361

52. Deuis, J.R. et al. (2017) Methods Used to Evaluate Pain Behaviors in Rodents. Frontiers in Molecular Neuroscience Volume 10–2017. 10.3389/fnmol.2017.00284

53. Dixon, W.J. (1980) Efficient Analysis of Experimental Observations. Annual Review of Pharmacology and Toxicology 20, 441–462. 10.1146/annurev.pa.20.040180.002301

